# Peripheral Nerve Transection Predominantly Drives Sympathetic Nerve Sprouting in Mouse Dorsal Root Ganglia

**DOI:** 10.1101/2025.11.07.687300

**Authors:** Sang Wook Shim, Hyoung Woo Kim, Yoon Kyung Lee, Clifford J. Woolf, Kihwan Lee, Seog Bae Oh

## Abstract

Sympathetic sprouting in dorsal root ganglia (DRG) is a feature of sympathetically maintained pain (SMP) following peripheral nerve injury, yet the factors determining its occurrence remain unclear. Here, we compare transection and crush injury models to determine if injury type or site influence sympathetic remodeling and pain. Using TH-IR immunostaining and Phox2b reporter mice to selectively label sympathetic fibers, we found that an L5 spinal nerve transection (SpNT) triggered robust sympathetic fiber sprouting and elevated norepinephrine (NE) levels in the DRG, correlating with a mechanical hypersensitivity reversed by chemical sympathectomy. In contrast, a partial sciatic nerve crush injury (PCI) produced long-lasting mechanical hypersensitivity without sympathetic sprouting or NE elevation and was unaffected by sympathectomy. Importantly, sympathetic sprouting was consistently more pronounced after transection injuries at both spinal and sciatic nerve sites, suggesting that injury type, rather than location, is a dominant factor shaping sympathetic remodeling. These findings establish nerve transection as a key driver of sympathetic sprouting and SMP, whereas crush-induced pain likely involves distinct non-sympathetic mechanisms. This distinction has important implications for pain subtype identification and treatment strategies.

## 1. Introduction

Chronic pain is a debilitating and complex condition affecting millions of people worldwide, with substantial variability in the underlying mechanisms across different individuals. Among these, sympathetically maintained pain (SMP) represents a subtype which arises from hyperactivity of the sympathetic nervous system. Despite clinical recognition of SMP in disorders such as complex regional pain syndrome (CRPS) and certain neuropathic pain states [4; 5; 16; 48; 70], the factors that lead to its occurrence in some individuals remain poorly understood.

Sympathetic sprouting into the dorsal root ganglion (DRG) has been widely documented in preclinical models of neuropathic pain, particularly those involving a nerve transection or ligation, such as chronic constriction injury (CCI) [31; 47], spinal nerve ligation (SpNL) [31; 62], partial sciatic nerve ligation (PSNL) [31; 53], and spared nerve injury (SNI) [45]. These models recapitulate neuropathic pain symptoms similar to those observed in humans [52], and are associated with sympathetic nerve sprouting [15; 39; 49; 50; 52] and increased levels of norepinephrine (NE) in the damaged DRG [55; 74], believed to be released from the sympathetic nerve terminals [56].

While all these models produce neuropathic pain-like behavioral changes, not all nerve injuries produce the same degree of aberrant sympathetic activity. The SpNL (or transection; SpNT) model [40; 41] and the SNI [45; 74] model show clear phenotypes of sympathetically maintained pain, but the CCI [3] and the PSNL [33; 53] models yield more variable results [31; 33]. These neuropathic pain models can be distinguished based on the surgical site and injury type [28], suggesting that the nature of nerve injury – specifically, the type and the location of the injury – may influence sympathetic remodeling and its contribution to pain [23; 28; 54].

Recent studies suggest that sciatic nerve crush injury models, particularly a partial sciatic nerve crush injury (PCI) model [9; 22; 29; 30] can be used as a model of neuropathic pain due to its ability to induce long-lasting mechanical hypersensitivity while preserving some axonal continuity and enabling regeneration [9; 29; 30]. Unlike other neuropathic pain paradigms [2; 10; 21], PCI applies a single crush across the sciatic trunk, resulting in a transient sensory loss followed by an onset of mechanical hypersensitivity. However, whether sympathetic activity contributes to the chronic pain in the PCI model has not been explored.

In this study, we investigate sympathetic activity in the PCI model and evaluate its relevance for studying the nature of SMP. We have compared the PCI model with the well-established SpNT model [18; 40]. We also compared the extent of sympathetic nerve sprouting in the DRG following crush and transection injuries at two distinct sites – the spinal nerve and the sciatic nerve. Our findings suggest that tactile hypersensitivity in the PCI model does not depend on sympathetic activity, whereas sympathetic nerve sprouting in the DRG is observed after a transection injury regardless of the location of the nerve injury. These findings provide novel insight into the role of different types of nerve injury in the development of SMP, which increases our understanding of SMP, and has the potential to select different treatment options for patients suffering from chronic pain following peripheral nerve injury.

## 2. Materials and Methods

### 2.1 Animals

Animal experiments were performed with 5- to 10-week-old both sexes C57BL/6J wild-type (WT) mice (Doo Yeol Biotech, South Korea) and *Phox2b-cre;tdTomato* mice. *Phox2b-cre;tdTomato* mice were generated by crossing hemizygous *Phox2b-cre* (strain # 016223) and homozygous Ai14 strain B6.*Rosa26-stop* (*flox*)-*tdTomato* (strain # 007914) which were obtained from the Jackson Laboratory (JAX). Mice were maintained in constant temperature (22 ± 1 ℃), humidity (55%), and a 12-hour light/dark cycle environment with standard laboratory chow and water available *ad libitum*. Handling and care of animals was performed following the Guideline for Animal Experiments, 2000, edited by the Korean Academy of Medical Sciences, which is consistent with the National Institutes of Health Guidelines for the Care and Use of Laboratory Animals, revised 1996, and were conducted according to institutional animal care and safety guidelines at Boston Children’s Hospital and Harvard Medical School. Experimental procedures were reviewed and approved by the Institutional Animal Care and Use Committee (IACUC) at Seoul National University (protocol code: SNU-200605-4-2, SNU-220111-3-1) and Boston Children’s Hospital (protocol code: 00002403).

### 2.2 Surgery

L5 spinal nerve transection (L5 SpNT) was performed as a modified version of Kim and Chung model [21; 27]. Mice were placed under isoflurane inhalation, the dorsal lumbar region was shaved, and treated with an iodine solution (Potadine). A unilateral incision was made parallel to the L6 vertebrate and the L6 transverse process was removed using blunt forceps. The L5 spinal nerve, which runs immediately below the L6 process, was isolated from connective tissue, and cut with fine spring scissors (15003-08, Fine Science Tools, Germany); 1 mm of the nerve was removed to prevent nerve regeneration. L4 spinal nerve crush (L4 SpNC) and transection (L4 SpNT) were performed similarly to L5 SpNT to expose the L4 spinal nerve. L4 spinal nerve was crushed for 15 s using Dumont SS fine forceps (11200-33, Fine Science Tools, Germany) for L4 SpNC and transected using fine spring scissors for L4 SpNT. Control mice were subjected to sham operation with the same surgical procedure without transection. The wound was irrigated with sterile saline and closed in two layers with 6-0 silk sutures (Ailee, Korea) and 9 mm skin clips (MikRon Precision, CA, USA). Partial sciatic nerve crush injury (PCI) was performed as previously described [29]. The sciatic nerve was crushed for 15s using a customized ultra-fine hemostat (13020-12, Fine Science Tools, Germany) with 30 μm gap. Sciatic nerve transection (ScNT) was performed in the same injury site with PCI using fine spring scissors. 2 mm piece of the sciatic nerve was removed to prevent the regeneration. The wound was closed in a suture of the overlying muscle facia and then the skin was closed with a skin clip. Mice were recovered in a warm, darkened cage and monitored until regaining consciousness and locomotion. In our study, we used a modified Chung model [21] by applying transection of L5 spinal nerve [27; 37; 44] to minimize the severe motor impairment observed with L4 SpNT (e.g., dragging hindlimb) [32], thereby enabling reliable behavioral assessment. To allow level-matched comparisons of sympathetic sprouting by injury type (transection *vs.* crush) and injury level (L4 *vs.* L5), we also included L4 SpNT in a subset of experiments.

### 2.3 Chemical sympathectomy

6-OHDA (162957, Merck, NJ, USA) was dissolved in sterile saline containing 0.01% (w/v) ascorbic acid (vehicle) and was injected intraperitoneally at a concentration of 200 mg/kg [34; 36; 40]. Control mice received an equivalent volume of vehicle alone. A total three injections were given every other day. The effect of chemical sympathectomy was validated using WB and ELISA.

### 2.4 von Frey test

To assess mechanical hypersensitivity, 50% paw withdrawal threshold was measured using *von Frey* filaments (North Coast Medical, CA, USA). Mice were acclimated in the cage for at least a week and then adapted in an acrylic cylinder (6.5 cm diameter, 17 cm height) on the metal mesh floor before the experiment at least three times. All animals were left in an acrylic cylinder on the metal mesh floor for an hour before the mechanical test. The 50% paw withdrawal threshold was determined based on the up-down method with an ascending series of *von Frey* filaments.

### 2.5 Sample preparation

#### 2.5.1 Sample preparation for immunohistochemistry

Mice were terminally anesthetized by intraperitoneal injection pf sodium pentobarbital (200 mg/kg) and perfused with 1X PBS followed by 4% PFA. Lumbar 4, 5 DRG were dissected and post-fixed overnight at 4 ℃ followed by cryopreservation in 30% sucrose in PBS for more than 3 days at 4 ℃. DRG tissues were embedded in OCT compound and cut into 10 μm sections with cryostat and directly mounted to Superfrost slides.

#### 2.5.2 Sample preparation for Western blot and ELISA

Mice were terminally anesthetized by intraperitoneal injection pf sodium pentobarbital (200 mg/kg) and perfused with PBS. L3-5 DRG and spleen were rapidly dissected, snap-frozen in liquid nitrogen, and stored at −80 ℃.

### 2.6 Tissue clearing

Whole DRG tissue clearing was performed using the Binaree Brain Clearing kit (Cat no. SHBC-001, Binaree, Korea). Briefly, fresh DRG tissue was dissected and immersed in 4% PFA for 24 h. Next day, the fixed tissue was directly moved to the tissue-clearing solution and incubated at 37 ℃ for 18 h on the shaker and then rinsed with the washing solution.

### 2.7 Immunohistochemistry

After washing in PBS, tissues were blocked and permeabilized in 5% normal donkey serum (NDS, Jackson ImmunoResearch, PA, USA) and 0.3% Triton X-100 in PBS for 1 h at RT. Primary antibodies were applied in 1% NDS, 0.3% Triton X-100 in PBS and incubated overnight at 4 ℃ in a humidified chamber. Fluorescence-conjugated secondary antibodies were incubated in 1% NDS, 0.3% Triton X-100 in PBS for 1 h at RT. Then images were obtained with a Zeiss LSM700/LSM980 confocal microscope.

For whole-mount staining of cleared L4 DRG, samples were blocked and permeabilized in 10% Bovine serum albumin (BSA) and 1% Triton X-100 in PBS for 2 h at RT. Primary antibodies were applied in 0.3% Triton X-100 in PBS and incubated 2 days at 4 ℃ on the shaker. After washing the sample with PBS, fluorescence-conjugated secondary antibodies were incubated in 0.3% Triton X-100 in PBS for 2 days at 4 ℃ on the shaker. Samples were incubated in mounting and storage solution for 24 h (Cat no. SHMS-050,

Binaree, Republic of Korea) after finishing staining. Then images were obtained with a Zeiss LSM980 confocal microscope.

### 2.8 Image analysis

#### 2.8.1 Quantification of sympathetic sprouting on 2D image

Five sections per sample were imaged for the analysis. The total area of TH-IR or Phox2b+ fibers, but not TH+ cell body, within the NeuN+ area (soma area) was measured using ImageJ software (NIH). The innervation density of the TH-IR fibers was calculated by dividing the area of TH-IR or Phox2b+ fibers (nm^2^) by NeuN+ area (μm^2^). To analyze Phox2b-negative and TH-IR fibers, the ROI of Phox2b+ regions were cleared from the TH channel in Image J. Subsequently, particle analysis of Phox2b- and TH-IR signals was performed to measure the innervation density (**Supplemental Fig. 1**).

#### 2.8.2 Quantification of sympathetic sprouting on 3D image

Five random images were extracted from z-stacked images of whole-mounted DRG. Z-stacks of images were processed and reconstructed in three dimensions with IMARIS software for representative 3D video images. The total area of TH-IR fibers, but not TH+ cell body, within the NeuN+ area (soma area) was measured using ImageJ software (NIH). The innervation density of the TH-IR fibers was calculated by dividing the area of TH-IR fibers (nm^2^) by the NeuN+ area (um^2^).

#### 2.8.3 Quantification of Iba1+ cell immunoreactivity in 2D images

Five sections per sample were imaged for analysis. The intensity and number of Iba1-positive cells within the NeuN+ area (soma rea) were measured using ImageJ software (NIH). Signal intensity was normalized to the NeuN+ area (μm^2^).

### 2.9 ELISA

#### 2.9.1 Quantification of NE level

The NE concentration in the spleen and DRG tissue was measured with a high-sensitivity ELISA kit (NOU39-K01, Eagle Biosciences, NH, USA) following the manufacturer’s manual. Three individual mice were pooled to measure NE levels in L3-5 DRG tissue. Tissues were sonicated (3-4 times, 10 s, 60% amplitude) in RIPA buffer (20-188, Merck, NJ, USA) containing protease inhibitor cocktail (P8340, Merck, NJ, USA) and phosphatase inhibitor cocktail (P3200, GenDepot, TX, USA). Protein content was determined by colorimetric assay (5000113, 5000114, BioRad, CA, USA).

### 2.10 Western blot

Frozen tissues were homogenized in RIPA lysis buffer (20-188, Merck, NJ, USA) containing protease inhibitor cocktail (P8340, Merck, NJ, USA) and phosphatase inhibitor cocktail (P3200, Gendepot TX, USA). After homogenizing the sample in a Minilys bead homogenizer (Precellys, Bertin, France) with zirconium beads (Precellys, Bertin, France), lysates were centrifuged at 10,000 rcf for 10 min. Protein concentration was determined by colorimetric assay (5000113, 5000114, BioRad, CA, USA).

Equal amounts of protein (40 μg) and protein size markers were separated by SDS-polyacrylamide gel electrophoresis (5% stacking gel, 10% resolving gel) followed by transfer to a PVDF membrane. Membranes were blocked in a 5% skimmed milk-containing Tris-buffered saline and 0.1% Tween-20 (TBS-T) at room temperature for 1 h before antibody incubation. Antibodies were diluted with blocking buffer and washed with 0.1% TBS-T after incubation. Primary antibodies were treated overnight at 4 ℃ shaker, and HRP-conjugated secondary antibodies were treated for 1 h at RT. Rabbit anti-tyrosine hydroxylase (BML-SA497-0100, Enzo Life Sciences, NY, USA) and mouse anti-β-actin (A5441, Sigma-Aldrich, MO, USA) were used at 1:1K and 1:10K respectively.

### 2.11 Statistical analysis

All statistical analyses were performed using IBM SPSS Version 26.0. The normal distribution of the data was analyzed with a Shapiro-Wilk test before further applying parametric or non-parametric statistical analyses. Unpaired Student’s *t*-test was used for normally distributed data. When data did not follow a normal distribution, the Mann-Whitney *U* test was used to compare the data between groups, and the Friedman test was used to formally detect differences between groups across repeated measures. Data were also tested for homogeneity of variance using Levene’s test. Investigators performing the behavioral test, quantitative histological staining and morphometric analyses were blinded either to the surgery or treatment group. Treatments were assigned to littermates at random by an independent observer. Data are presented as mean ± standard error of the mean (SEM). *p* < 0.05 was considered significant.

## 3. Results

### Time course of mechanical hypersensitivity following spinal transection and sciatic partial crush injuries

We assessed mechanical hypersensitivity in two neuropathic pain models: L5 spinal nerve transection (SpNT) and partial sciatic nerve crush injury (PCI) (**Fig. 1A**). L5 SpNT produced mechanical hypersensitivity by day 7 that persisted through day 30 (**Fig. 1B**). PCI showed a delayed onset of hypersensitivity, first emerging at day 15, and also persisting to day 30 (**Fig. 1C**), which confirms previous reports [9; 29; 30]. Unlike selective branch/root models like the spared nerve injury [10] or L5 spinal nerve crush models [60], which spare an entire branch or root, the PCI model [9; 29; 30] applies a single crush across the sciatic trunk, injuring all fascicles while variably sparing a subset of axons within them, leading to distinct temporal dynamics of sensory changes, including delayed onset of mechanical hypersensitivity. Therefore, while both models result in persistent mechanical hypersensitivity, the timing of onset differs.

**Figure 1.**
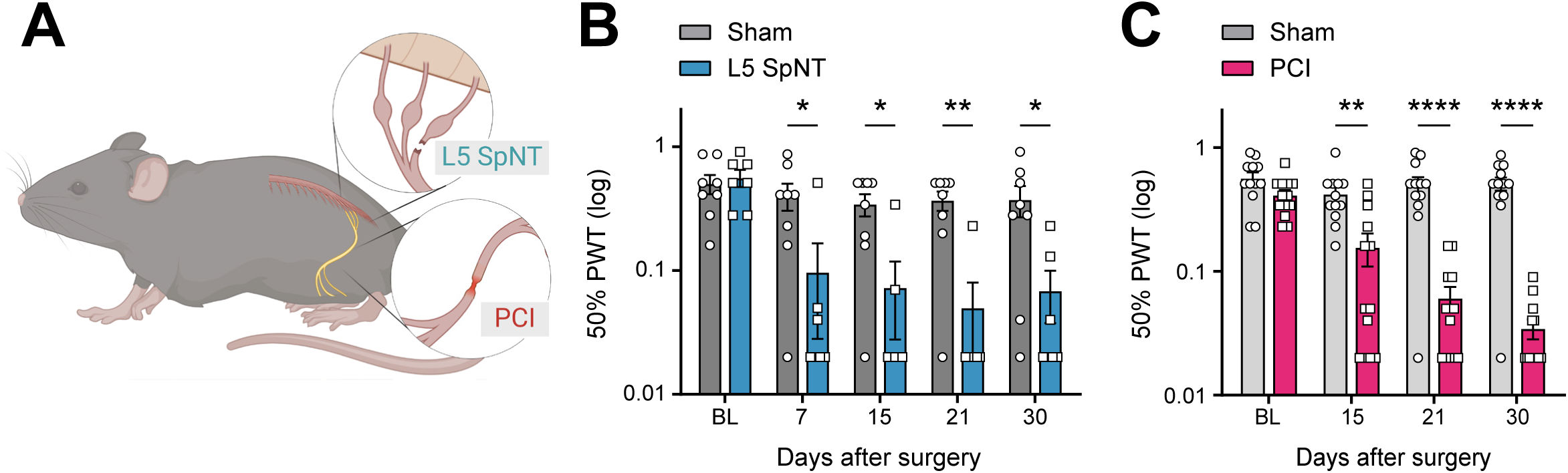
Tactile hypersensitivity following spinal nerve transection and partial sciatic nerve crush injury models. **(A)** Schematic image of L5 SpNT and PCI in mice. **(B)** Mechanical hypersensitivity assessed by 50% paw withdrawal thresholds of ipsilateral hind paws at 7, 15, and 30 days following L5 SpNT. **(C)** 50% paw withdrawal thresholds measured in the ipsilateral hind paw at 15, 21 and 30 days following PCI. Data represent mean ± SEM (n=12–13 mice per group); \**p* < 0.05, \*\**p* < 0.01, \*\*\*\**p* < 0.0001, Mann–Whitney *U* test. L5 SpNT, L5 spinal nerve transection; PCI, Partial sciatic nerve crush injury

### TH-IR fiber sprouting and norepinephrine elevation are exclusive to SpNT

Previous studies showed that sympathetic fiber sprouting occurs in the L5 DRG after a L5 SpNT [6; 40], and an increase of norepinephrine (NE) levels in the DRG has been documented in various neuropathic pain models [74]. NE is released from sympathetic nerve terminals and can sensitize sensory neurons [26; 55; 61; 66; 74]. Tyrosine hydroxylase (TH) is an enzyme involved in dopamine and norepinephrine synthesis and its immunoreactivity can be used as an indicator of sympathetic neurons [8].

To determine whether sympathetic activation underlies the hypersensitivity observed in these two models, we first examined TH immunoreactivity (TH-IR) and NE levels in the DRG. Following L5 SpNT, the density of TH-IR fibers markedly increased in the ipsilateral L5 DRG at days 7 and 15, but returned to baseline by day 30 (**Figs. 2A and B**). By contrast, TH-IR fiber density in the L4 DRG remained unchanged following a PCI at all time points (**Figs. 2A and C**), as well as in sham-operated mice (**Supplemental Fig. 2**). Furthermore, ipsilateral L4 DRGs from L5 SpNT mice did not increase TH-IR fiber innervation at days 15 (**Supplemental Fig. 3**), indicating that the sympathetic sprouting occurs exclusively in the injured DRG following SpNT.

**Figure 2.**
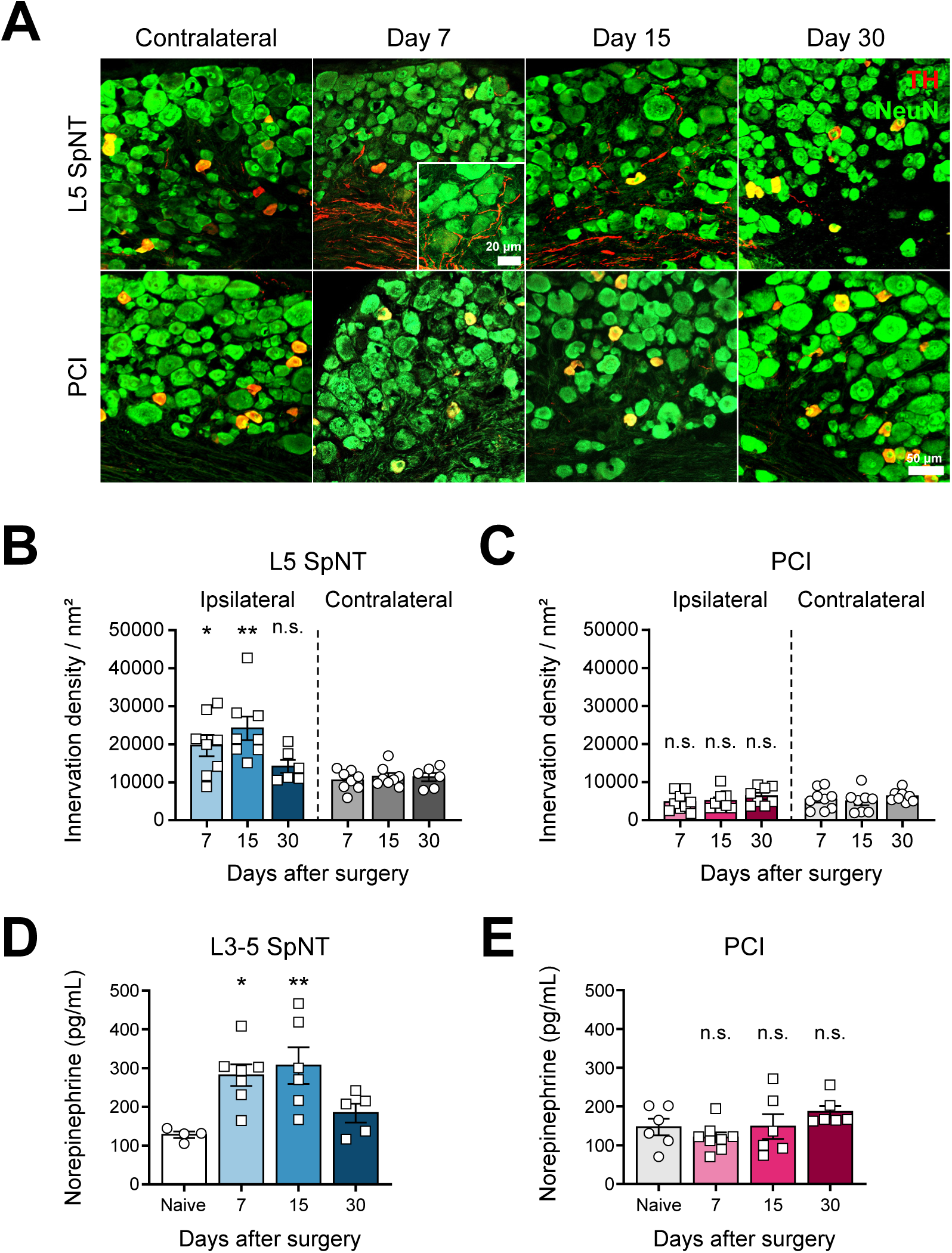
TH-IR fiber sprouting occurs in the L5 DRG after SpNT but not PCI. **(A)** Representative immunostaining images of TH-immunoreactive (TH-IR) fibers (nm^2^) (red) in NeuN+ DRG neurons (μm^2^) (green) at 7, 15, and 30 days after L5 SpNT and PCI. **(B)** Quantitative analysis of TH-immunoreactive (TH-IR) fibers (nm^2^) (red) in NeuN+ DRG neurons (μm^2^) (green) at 7, 15, and 30 days after L5 SpNT. **(C)** Quantitative analysis in the L4 DRG at corresponding time points after PCI showed no significant increase in TH-IR fiber density. Data represent mean ± SEM (n=4–5 mice per group, average of 5 sections per mouse); \**p* < 0.05, \*\**p* < 0.01, two-tailed unpaired *t*-test. **(D, E)** Quantification of NE levels in L3–L5 DRG at indicated time points following L3–L5 SpNT and PCI. Significant NE upregulation occurred only after SpNT. Data represent mean ± SEM (n=4–8 mice per group and time point; pooled samples of three mice per data point). \**p* < 0.05, \*\**p* < 0.01, two-tailed unpaired *t*-test. TH, Tyrosine hydroxylase; IR, Immunoreactive; DRG, Dorsal root ganglia; L5 SpNT, L5 spinal nerve transection; PCI, Partial sciatic nerve crush injury; NE, Norepinephrine

Correspondingly, NE levels in the ipsilateral L3-5 DRG were significantly elevated after L3-5 SpNT on days 7 and 15, but not at day 30 (**Fig. 2D**), while there was no elevation in the PCI model (**Fig. 2E**). Together, these results indicate that a SpNT injury, not a PCI, triggers sympathetic remodeling in the DRG.

### Injury type—not location—determines the extent of sympathetic sprouting

The L5 SpNT and PCI models differ not only in the type but also in the location of the nerve injury (**Supplemental Fig. 4**). Therefore, we expanded our analysis to include a crush injury of the L4 spinal nerve (L4 SpNC) and sciatic nerve transection (ScNT), enabling matched comparisons of sympathetic sprouting by injury type (transection *vs.* crush) and injury level (L4 *vs.* L5). The innervation density of TH-IR fibers in the L4 DRG was assessed by whole-mount immunohistochemistry in all four models. Sympathetic nerve sprouting was not observed in the L4 DRG after an L4 SpNC (**Supplemental Figs. 5A-C**), but it increased after a ScNT (**Supplemental Figs. 5D-F**, **Supplemental Movie 1**). Results for L4 SpNT and PCI were consistent with the findings presented in **Figure 2**.

Since TH is also a marker of c-LTMR sensory neurons [35], we employed Phox2b-Cre;tdTomato mice to verify that the TH-IR fibers are indeed of sympathetic origin [20]. Following L4 spinal nerve injury, both transection (SpNT) (**Figs. 3A and B**) and crush (SpNC) (**Figs. 3A and C**) showed increased Phox2b+ fiber innervation in DRG, with a significantly greater increase after transection. Importantly, TH-IR but Phox2b-negative areas did not increase (**Supplemental Fig. 6**), indicating that the signal increase is primarily sympathetic. The small Phox2b+ increase after an L4 SpNC despite no detectable TH-IR change is consistent with marker sensitivity/contrast—Phox2b provides a higher specificity for sympathetic lineage neurons, whereas TH-IR includes baseline c-LTMR signals that can mask subtle gains. In contrast, after a sciatic nerve injury, only a transection (ScNT) injury increased Phox2b+ fiber innervation in DRG (**Figs. 3D, E**), whereas PCI showed no significant increase (**Figs. 3D, F**). These level-matched comparisons show that nerve transection elicits a robust sympathetic sprouting response, crush injuries at most only a limited increase, and injury type—rather than anatomical site—is the principal determinant of sympathetic sprouting.

**Figure 3.**
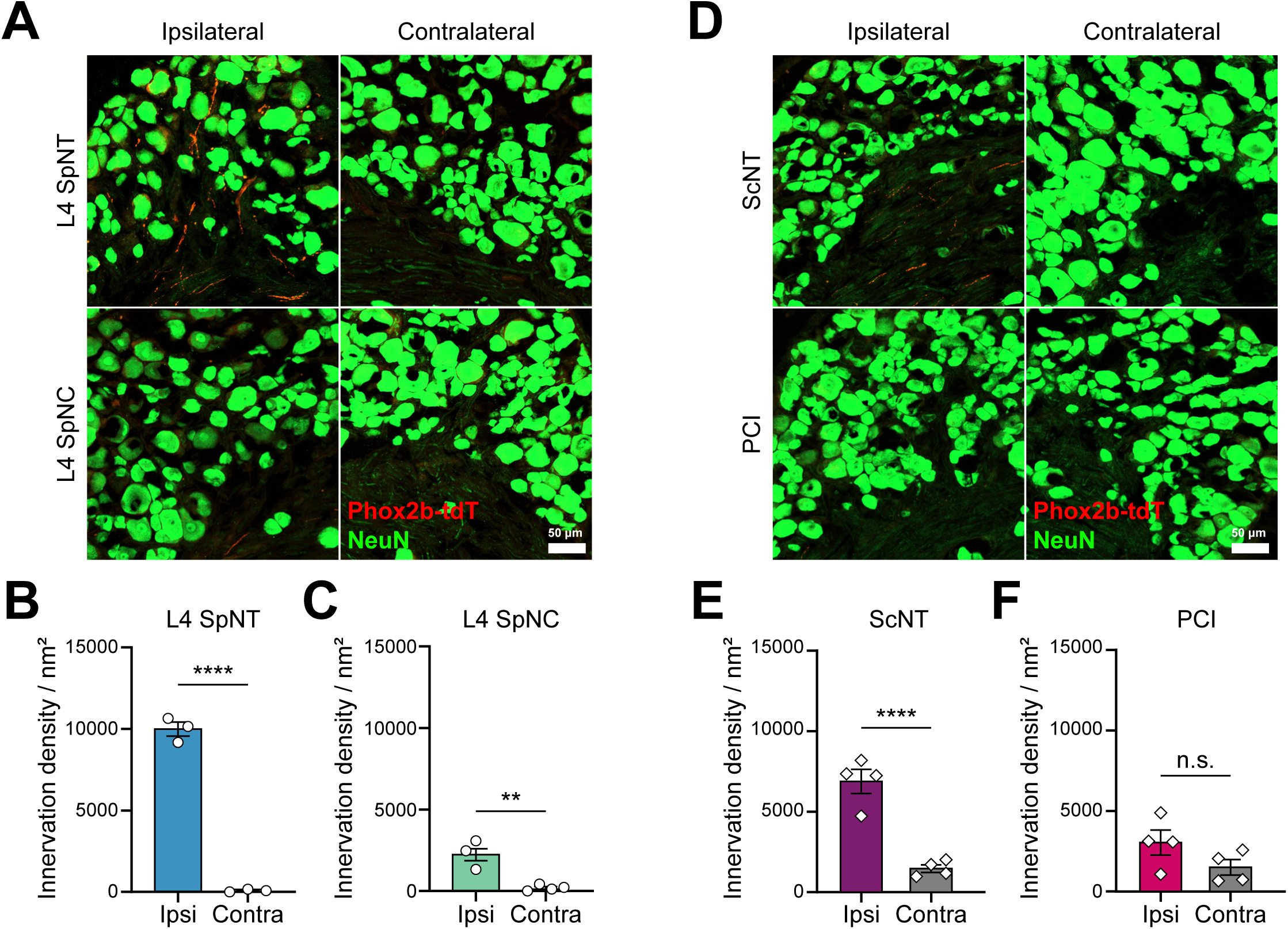
Genetic labeling with Phox2b-tdTomato confirms that TH-IR fiber sprouting depends on injury type rather than injury location. **(A)** Representative fluorescent images of sympathetic fiber sprouting (Phox2b-tdTomato fibers, red) around NeuN+ neurons (green) in L4 DRG at 15 days post-L4 SpNT and L4 SpNC. **(B)** Quantitative analysis of Phox2b+ fibers revealed significant increases in tdTomato+ fiber density following L4 SpNT compared to control group (n=3 mice per group). **(C)** Similar analysis following L4 SpNC, showing significant tdTomato+ sympathetic sprouting (n=3 mice per group). **(D)** Representative fluorescent images of sympathetic fiber sprouting (Phox2b-tdTomato fibers, red) around NeuN+ neurons (green) in L4 DRG at 30 days post-ScNT and PCI. **(E)** Quantitative analysis of Phox2b+ fibers revealed significant increases in tdTomato+ fiber density following ScNT compared to control group (n=4 mice per group). **(F)** Similar analysis following PCI, showing significant tdTomato+ sympathetic sprouting (n=4 mice per group). Data represent mean ± SEM. \*\**p* < 0.01, \*\*\*\**p* < 0.0001, two-tailed unpaired *t*-test. L4 SpNT, L4 spinal nerve transection; L4 SpNC, L4 spinal nerve crush; ScNT, Sciatic nerve transection; PCI, Partial sciatic nerve crush injury

### Sympathectomy attenuates mechanical hypersensitivity by SpNT, but not by PCI

To assess the functional contribution of sympathetic activity to pain-related behaviors, we employed a chemical sympathetic block prior to the injuries through systemic injection of 6-hydroxydopamine (6-OHDA; total 600 mg/kg, *i.p.*) (**Fig. 4A and Supplemental Fig. 7A**), which elevates the production of reactive oxygen species in catecholaminergic neurons, leading to their death [51]. Systemic administration of 6-OHDA significantly reduces both TH expression (**Supplemental Figs. 7B and C**) and NE (**Supplemental Fig. 7D**) in the spleen, and the reduction persisted for up to 30 days (**Fig. 4B**), indicating that the effects of 6-OHDA were sustained throughout the experiment.

**Figure 4.**
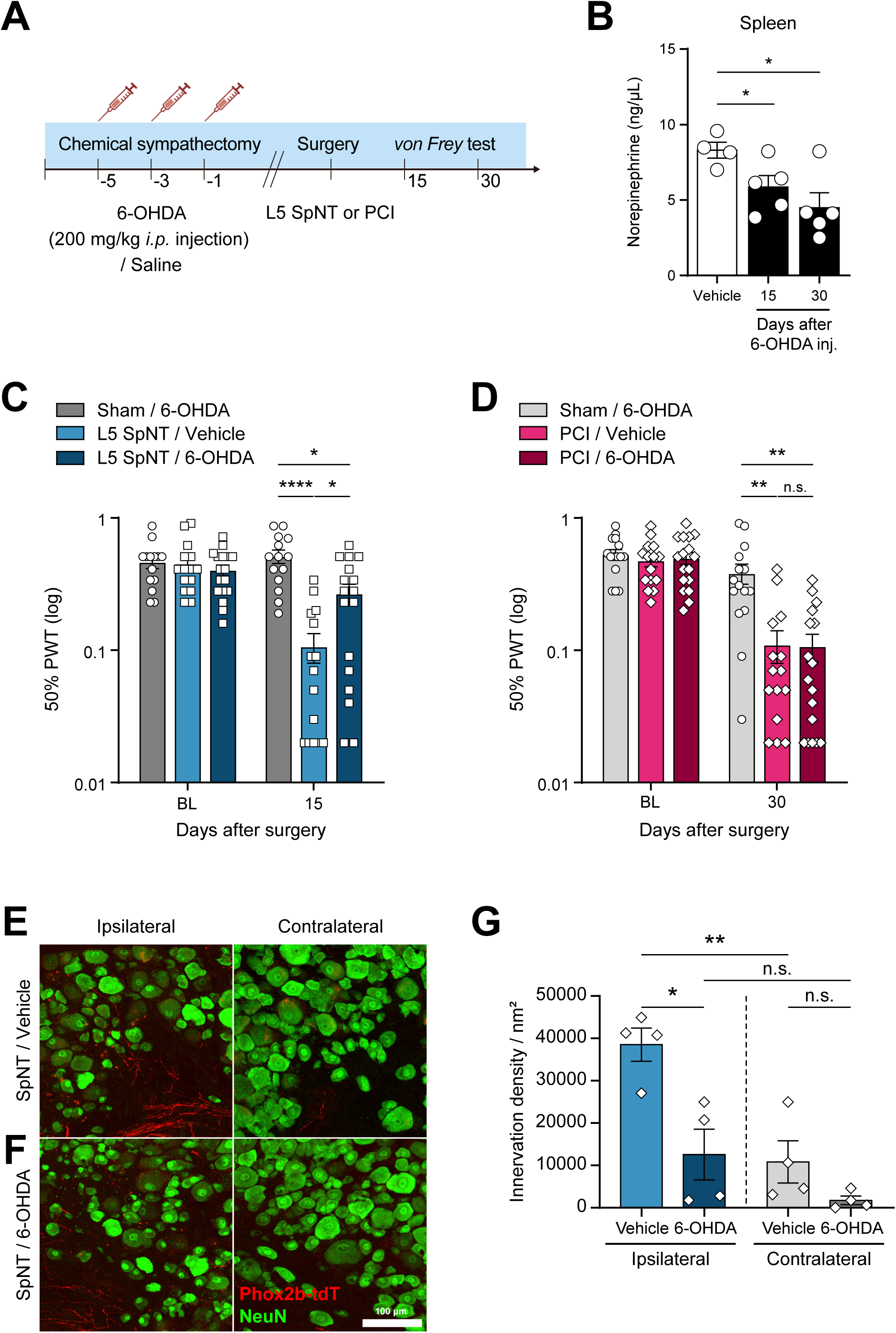
Effect of chemical sympathectomy on mechanical hypersensitivity. **(A)** Experimental timeline indicating the schedule of chemical sympathectomy and nerve injury. **(B)** Validation of sympathectomy efficacy at 15 and 30 days post-injection. Data represent mean ± SEM (n=4– 5 mice per group). \**p* < 0.05, two-tailed unpaired *t*-test. **(C)** 50% paw withdrawal thresholds of ipsilateral hind paw in 6-OHDA- *vs.* vehicle-treated mice 15 days after L5 SpNT. Mechanical allodynia was partially alleviated by chemical sympathectomy (n=14–17 mice per group). **(D)** No significant alleviation of mechanical allodynia was observed following sympathectomy in the PCI model (n=15–17 mice per group). Data represent mean ± SEM. \**p* < 0.05, \*\**p* < 0.01, \*\*\*\**p* < 0.0001, Mann–Whitney *U* test. **(E-G)** Representative immunostaining images and quantitative analysis of Phox2b+ fibers (nm2) (red) in NeuN+ DRG neurons (μm2) (green) in 6-OHDA *vs.* vehicle mice 15 days after L4 SpNT. Data represent mean ± SEM (n=4 mice per group, average of 5 sections per mouse). Two-tailed unpaired *t*-test. 6-OHDA, 6-hydroxydopamine; *i.p.*, intraperitoneal; L5 SpNT, L5 spinal nerve transection; PCI, Partial sciatic nerve crush injury

Based on the onset of hypersensitivity (**Fig. 1**) and sympathetic nerve sprouting (**Fig. 2-3**), we measured the effect of chemical sympathectomy on day 15 in the L5 SpNT mice and day 30 in the PCI model. 6-OHDA injection significantly attenuated the development of mechanical hypersensitivity in the L5 SpNT model (**Fig. 4C**), as previously reported [40; 41], but had no effect in PCI (**Fig. 4D**), supporting that sympathetic activity contributes to pain after SpNT but not PCI. Consistent with the analgesic efficacy, 6-OHDA reduced TH-IR (**Supplemental Figs. 8A, B**) and Phox2b+ fiber densities in injured DRG 15 days after SpNT (**Figs. 4E-G**), indicating functional contribution of sympathetic sprouting to pain behavior.

## 4. Discussion

In this study, we demonstrate that the PCI model does not exhibit characteristics of sympathetically maintained pain. By comparing the effects of transection and crush injuries at two distinct nerve locations, we investigated their relationship with sympathetic nerve sprouting in the DRG and associated pain phenotypes. Our findings reveal a strong relationship between transection (SpNT, ScNT) and the extent of sympathetic sprouting, with crush producing smaller (SpNC) or no (PCI) sprouting. Together with the selective efficacy of chemical sympathectomy in SpNT but not PCI, these findings define two mechanistic classes of neuropathic pain: sympathetically maintained (SpNT) and sympathetically independent (PCI).

To address what drives sympathetic sprouting, we compared the effect of nerve transection and crush injuries at two distinct anatomical locations (spinal nerve *vs.* sciatic nerve). Previous studies suggested that proximity of the injury site to the DRG correlates with the severity of DRG damage [19; 25; 63] and that sympathetic nerve sprouting is more pronounced when injury occurs near the DRG [28]. However, our findings indicate that the proximity alone is neither sufficient nor necessary: SpNC (proximal crush) produced only a limited Phox2b+ fiber increase, both SpNT (proximal transection) and ScNT (distal transection) elicited robust sprouting, and PCI (distal crush) showed no significant increase (**Fig. 3 and Supplemental Fig. 5**). Thus, the injury type—transection versus crush—is the primary determinant of sympathetic fiber sprouting, while location may primarily influence sprouting latency and, at most, magnitude [39; 45; 67]. Collectively, these comparisons indicate that the presence and extent of sympathetic nerve sprouting in the DRG depends predominantly on the type rather than the proximity of the nerve injury to the DRG.

Sympathetic nerve fibers play a critical role in pain generation following nerve injury, primarily through NE release at nerve terminals [46]. NE causes neuronal hyperexcitability by interacting with adrenergic receptors on sensory neurons [24; 69; 73]. An upregulation of alpha-1 and/or 2 adrenergic receptors on sensory neurons has been reported in nerve injury conditions [24; 62], which may increase neuronal excitability in response to NE. β3 adrenoceptors also play an important role in pain generation by upregulating ATP release in sensory neurons [26]. NE-evoked vasoconstriction may also indirectly drive sensory neuron firing [65]. Consistent with this, we observed NE upregulation in the DRG after SpNT (**Fig. 2**), and 6-OHDA both reduced DRG sympathetic sprouting and attenuated mechanical hypersensitivity (**Fig. 4**). These results support a contribution of sympathetic input to pain behavior after a transection injury and align with Zheng et al, who showed that NE from sprouted sympathetic fibers drives a synchronized DRG “cluster firing” that correlated with spontaneous pain-related behavior in the SNI model [15; 74].

We assessed sympathetic fiber sprouting in the DRG by using both TH-IR and Phox2b-tdTomato fiber density measurements in the soma area, which revealed a marker-specific discrepancy in the crush model. While TH-IR fiber sprouting increased only after transection injury (**Figs. 2A-C**), Phox2b+ fibers also increased in L4 SpNC, with a much greater effect in transection (**Figs. 3A-C**); by contrast, PCI showed no significant Phox2b+ increase at matched time points (**Figs. 3D-F**). This divergence is consistent with known limitations: TH labels both sympathetic fibers and c-LTMR neurons in the DRG, potentially masking subtle increases in sympathetic sprouting, whereas Phox2b is selective for sympathetic lineage but not expressed in all mature sympathetic neurons. To further clarify the TH signal, we analyzed TH+ DRG neurons and found that their numbers decrease following SpNT, making it unlikely that the increased TH-IR fibers originate from c-LTMR neurons (**Supplemental Fig. 9**). Moreover, the TH-IR+/Phox2b- area did not increase (**Supplemental Fig. 6**), indicating that the injury-induced TH-IR gain largely reflects sympathetic fibers rather than c-LTMR sensory fibers. Nonetheless, because TH can label c-LTMRs and Phox2b is not expressed in all mature sympathetic neurons, definitive attribution would be strengthened by retrograde tracing of DRG-projecting sympathetic neurons [6; 74]. Notably, an increased number of labeled sympathetic neurons has been reported in the L5 spinal nerve ligation model [6]. Consistent with these findings [31; 40], we observed robust increases in TH-IR and Phox2b+fibers following SpNT (**Fig. 2, 3**). Furthermore, NE levels were concomitantly elevated (**Fig. 2**), collectively supporting that the injury-induced TH-IR fibers predominantly originate from sympathetic neurons and actively release NE at their terminals.

Previous mechanistic studies have emphasized the role of neurotrophic factors, such as nerve growth factor (NGF) and brain-derived neurotrophic factor (BDNF), in driving sympathetic nerve sprouting within the DRG following nerve injury [11; 59]. These growth factors can originate from sensory neurons and non-neuronal satellite glial cells (SGCs), and both facilitate sympathetic fiber basket formation around sensory neurons [50; 58; 75]. Since the type and location of nerve injury could differentially influence neurotrophic factor release, future research should examine if non-neuronal cell populations, particularly SGCs, respond differently depending on the nature of the injury. Developmental pathways that guide sympathetic-sensory wiring may also be reactivated in adulthood. For instance, motor neurons and Schwann cell precursors (SCPs) regulate sympathetic fiber positioning during development, and their absence causes premature ingrowth into sensory ganglia [13]. Whether similar mechanisms are involved in injury-induced sprouting in adults remains to be determined, but differences in motor neuron or Schwann cell precursor responses may contribute to the distinct outcomes observed between crush and transection injuries.

Sympathetic nerve sprouting around sensory neurons following a peripheral nerve injury has long been considered the anatomical basis for SMP. However, its exact functional contribution to pain behaviors remains controversial. Spontaneous pain may originate from injured afferent neurons or hypersensitized intact afferents adjacent to injured neurons [7; 72], with spontaneous ectopic activity arising from DRG soma or neuromas [1; 7; 12]. Recent work in the SNI model, characterized by transection of two distal branches of the sciatic nerve, showed synchronized cluster firing in DRG neurons that correlated with spontaneous pain behaviors, driven by norepinephrine released from sympathetic fibers sprouted into the DRG [15; 74]. Taken together with our finding that transection (but not PCI) elicits DRG sympathetic sprouting and responds to sympathectomy, these observations suggest that sprouting is most likely to influence ongoing/spontaneous pain rather than evoked responses alone. Accordingly, beyond mechanical hypersensitivity, spontaneous pain behavior should be explicitly identified in each model to establish the functional significance of sympathetic sprouting.

Sympathetic sprouting can also occur in peripheral targets such as the upper dermis of the skin in some neuropathic pain conditions [17; 43; 71], raising the possibility that sympathetic fiber sprouting in tissues outside the DRG may also contribute to SMP phenotypes. Future work should test whether extra-ganglionic remodeling adds to behavioral outcomes in specific models.

It is important to acknowledge a limitation of the transection model. Removing a 1–2 mm nerve segment does not fully prevent axonal regeneration [68], rather, regenerating axons often fail to achieve precise target re-innervation. This may explain the decline in sympathetic sprouting and norepinephrine levels by day 30 after an L5 SpNT (**Fig. 2**). Partial bridging of the transection gap could allow reconnection between proximal and distal stumps, reducing denervation-driven cues for sympathetic ingrowth [57]. Alternatively, incomplete or misdirected reinnervation might alter trophic and immune signaling to the DRG, consequently impacting the maintenance of sympathetic fibers [6].

We also recognize the limitations of systemic 6-OHDA sympathectomy rather than localized microsympathectomy [64]. Systemic 6-OHDA administration is widely used to assess sympathetic involvement in pain [40; 41; 62], but can exert off-target, systemic immune effects [76], and does not isolate DRG-adjacent sympathetic inputs as precisely as a microsympathectomy. Nevertheless, we selected 6-OHDA because reliable surgical access to the relevant paravertebral sympathetic ganglia in our mouse cohorts is technically difficult precluding localized ablation. 6-OHDA reduced TH-IR/Phox2b+ fibers and lowered NE (trend) (**Supplemental Fig. 8C**) in DRG while attenuating SpNT hypersensitivity; meanwhile, IBA1 immunoreactivity in injured DRG was unchanged versus vehicle at day 15 (**Supplemental Fig. 10**), consistent with reports showing that 6-OHDA does not significantly alter DRG macrophage function [42]. Therefore, under our experimental conditions, the behavioral changes are better explained by a reduced sympathetic input rather than a reduction in DRG macrophages. Nevertheless, Zhu et al. [76] used microsympathectomy and observed immune changes, and 6-OHDA can modulate dendritic/T cell cytokines [42]—so a systemic sympathectomy may under- or over-estimate neuroimmune contributions. Future work using localized surgical microsympathectomy or targeted genetic strategies will be important to refine site specificity and causality.

Additionally, we did not detect significant sex-dependent effects on sympathetic sprouting, NE levels, or hypersensitivity across time points in either model (SpNT, PCI). However, sample sizes for each sex were insufficient for powered statistics, so sex-disaggregated results are not presented and pooled data are shown.

In conclusion, our results demonstrate that PCI-induced pain phenotypes appear sympathetically independent, in contrast to the sympathetically maintained state following nerve transection. These findings highlight the injury type as a critical determinant of sympathetic nerve sprouting in DRG and related mechanisms, with implications for identifying pain subtypes after traumatic nerve injury. Early identification of sympathetically dependent versus -independent pain could enable more targeted and effective therapies [14; 38] for patients with neuropathic pain.

## Acknowledgements

This research was supported by the National Research Foundation of Korea (NRF) grant funded by the Korean government (MSIT) RS-2021-NR059709, RS-2023-00264409, RS-2024-00441103 (to S. B. Oh) RS-2023-00272846 (to S. W. Shim) and RS-2023-00240792 (to H. W. Kim). We thank the Boston Children’s Hospital IDDRC Cellular Imaging Core, funded by NIH P50 HD105351. All authors gave their final approval and agreed to be accountable for all aspects of the work.

## Author contributions

Conceptualization: S. W. Shim, H. W. Kim and S. B. Oh; Methodology and investigation: S. W. Shim, H. W. Kim and Y. K. Lee. Resources : C. J. Woolf and S. B. Oh. Writing (original draft): S. W. Shim, H. W. Kim. Writing (editing): S. W. Shim, H. W. Kim, C. J. Woolf, K. Lee and S. B. Oh. Project supervision: K. Lee and S. B. Oh.

## Data availability

The authors confirm that the data supporting the findings of this study are available within the article and its supplementary materials. Additional study and data details are available from the corresponding author, [S. B. Oh], on special request.

## Declaration of interests

C.J.W. is a founder of Nocion Therapeutics, Quralis and BlackBox Bio and an SAB member of Lunbeck Pharma, Tafalgie Therapeutics, and Axonis. S.B.O. is a founder of OhLabBio.

## Notes

### Competing Interest Statement

The authors have declared no competing interest.

