## Supplementary Information (PDF) for "Peripheral Nerve Transection Predominantly Drives Sympathetic Nerve Sprouting in Mouse Dorsal Root Ganglia"

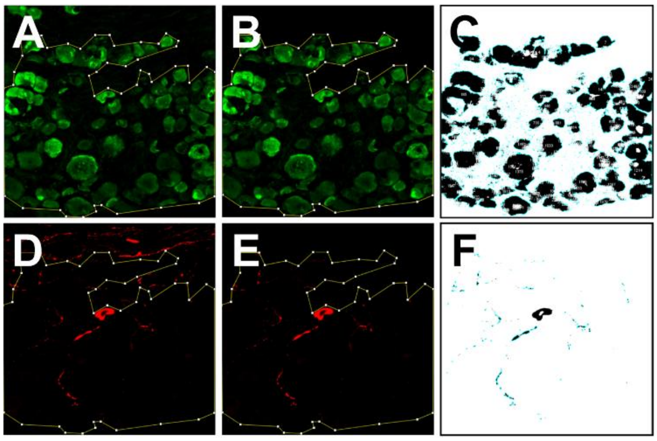


**Supplemental figure 1. Analysis methods of the innervation density of TH-IR fiber.**

**(A)** NeuN+ regions were manually delineated using ImageJ software. **(B)** Signals outside these regions were excluded using the "Clear Outside" plugin. **(C)** Particle analysis was performed for NeuN+ regions (green) after adjusting the threshold. **(D)** The identified NeuN+ region areas were applied to the TH+ channel (red). **(E)** Signals outside the NeuN+ regions were excluded using the "Clear Outside" plugin. **(F)** Particle analysis of TH-IR fibers was conducted after threshold adjustment, with a size range of 1-100 pixels$^2$ set to exclude signals from TH+ neurons. TH, Tyrosine hydroxylase; IR, Immunoreactive

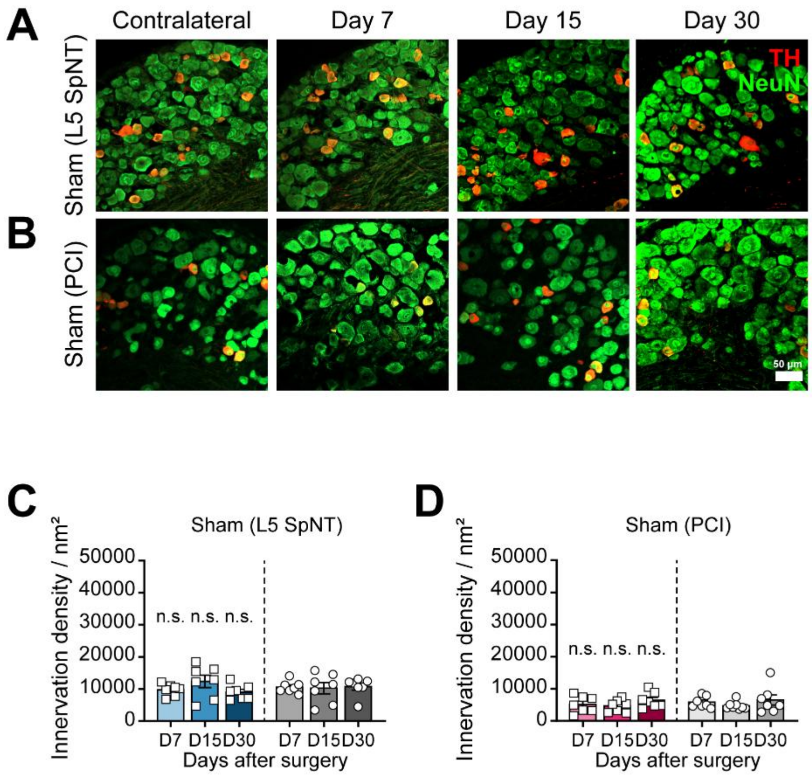


**Supplemental figure 2. TH-IR fibers are not sprouted into DRGs following sham surgery of L5 SpNT and PCI.**

**(A-D)** Representative immunostaining images and quantitative analysis of TH-IR fibers ($nm^2$) (red) in NeuN+ DRG neurons ($\mu m^2$) (green) at 7, 15, and 30 days after Sham surgery of L5 SpNT and PCI. Data represent mean ± SEM (n=7-8 mice per group, average of 5 sections per mouse); two-tailed unpaired *t*-test. TH, Tyrosine hydroxylase; IR, Immunoreactive; DRG, Dorsal root ganglia; L5 SpNT, L5 Spinal nerve transection; PCI, Partial sciatic nerve crush injury; n.s., not significant

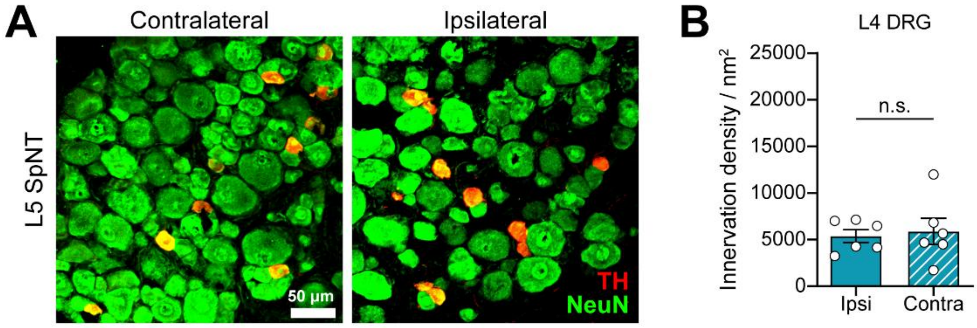


**Supplemental figure 3. TH-IR fibers are not sprouted into L4 DRG following L5 SpNT.**

**(A, B)** Representative immunostaining images and quantitative analysis of TH-IR fibers ($nm^2$) (red) in NeuN+ DRG neurons ($\mu m^2$) (green) in L4 DRG at 15 days after L5 SpNT. TH-IR fibers to ipsilateral L4 DRG were not observed after L5 SpNT. Data represent mean ± SEM (n=6 mice per group, average of 5 sections per mouse); Two-tailed unpaired *t*-test. TH, Tyrosine hydroxylase; IR, Immunoreactive; DRG, Dorsal root ganglia; L5 SpNT, L5 Spinal nerve transection; n.s., not significant

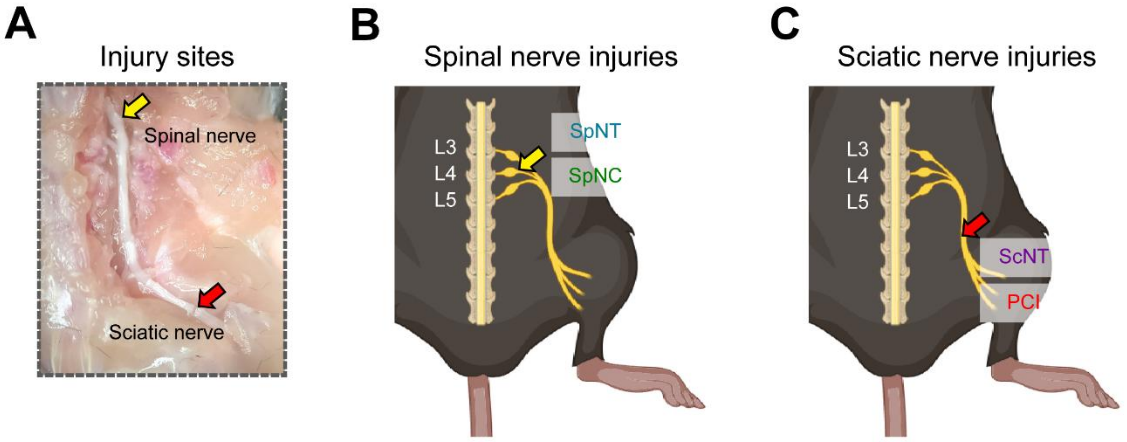


**Supplemental figure 4. Schematic images of L4 SpNT, L4 SpNC, ScNT and PCI indicate the variations in the location and type of nerve injury.**

**(A)** Injury sites of spinal nerve injuries (yellow arrow) and sciatic nerve injuries (red arrow) in WT mice. **(B)** Schematic image of L4 SpNT and L4 SpNC. **(C)** Schematic image of ScNT and PCI. WT, wild type; L4 SpNT, L4 Spinal nerve transection; L4 SpNC, L4 Spinal nerve crush; ScNT, Sciatic nerve transection; PCI, Partial sciatic nerve crush injury

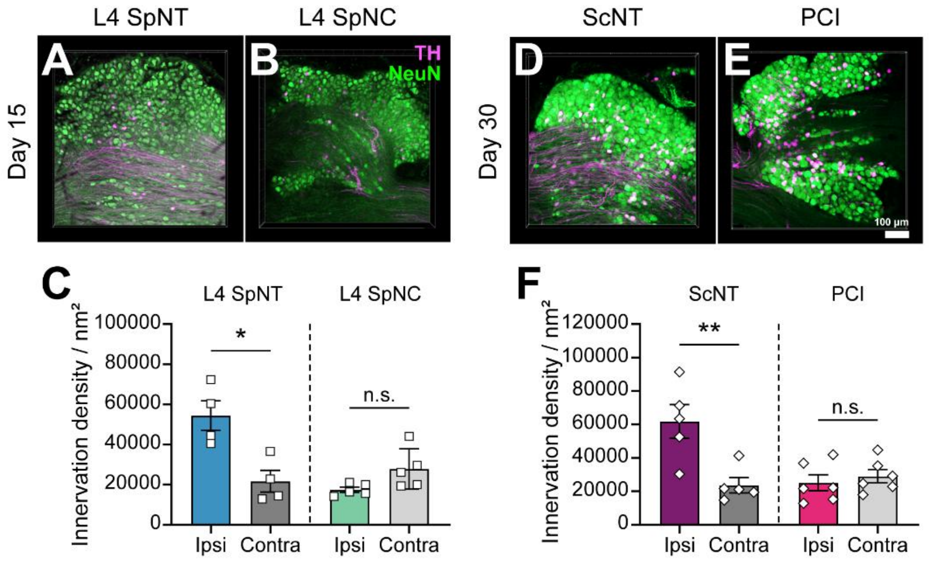


**Supplemental figure 5. TH-IR fiber sprouting depends on injury type rather than injury location.**

**(A-C)** Representative whole-mount immunostaining and quantification of TH-IR fibers (magenta) in NeuN+ DRG neurons (green) at 15 days following SpNC and L4 SpNT. Significant sympathetic fiber sprouting was observed only in the SpNT model (n=4–5 mice per group; average of 5 sections per mouse). **(D-F)** Whole-mount staining of L4 DRG following PCI and ScNT. Significant TH+ sympathetic sprouting occurred exclusively after transection injuries. (n=5–6 mice per group; average of 5 sections per mouse). Data represent mean ± SEM; *$p$ < 0.05, **$p$ < 0.01, two-tailed unpaired $t$-test. L4 SpNT, L4 Spinal nerve transection; L4 SpNC, L4 Spinal nerve crush; ScNT, Sciatic nerve transection; PCI, Partial sciatic nerve crush injury; n.s., not significant

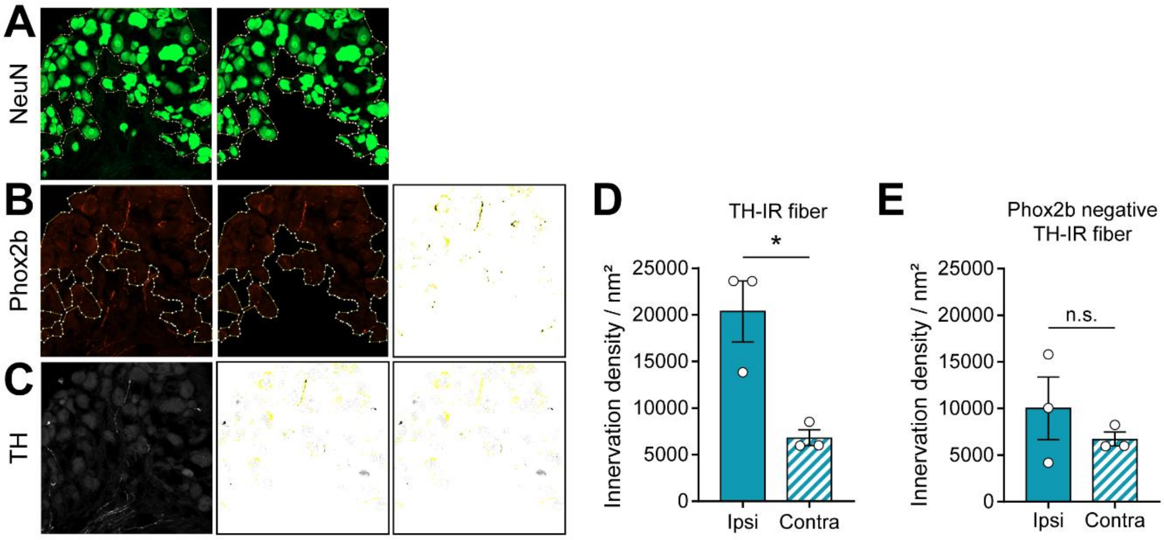


**Supplemental figure 6. Phox2b-negative and TH-IR fibers are not increased after L4 SpNT.**

**(A)** NeuN+ regions (green) were manually delineated using Image J software. **(B)** The area of Phox2b+ fibers (red) within the NeuN+ region was measured. **(C)** The ROI of Phox2b+ regions was excluded from TH-IR regions using the "Fill" plugin. **(D)** The innervation density of TH-IR fibers within the NeuN+ region was analyzed 15 days after L4 SpNT in Phox2b-tdTomato mice. Data represent mean ± SEM (n=3 mice per group, average of 5 sections per mouse); *$p < 0.05$, two-tailed unpaired *t*-test. **(E)** The innervation density of Phox2b- and TH+ fibers within the NeuN+ region was analyzed 15 days after L4 SpNT in Phox2b-tdTomato mice. Data represent mean ± SEM (n=3 mice per group, average of 5 sections per mouse); two-tailed unpaired *t*-test. ROI, Region of interest; TH, Tyrosine hydroxylase; IR, Immunoreactive; L4 SpNT, L4 Spinal nerve transection; n.s., not significant

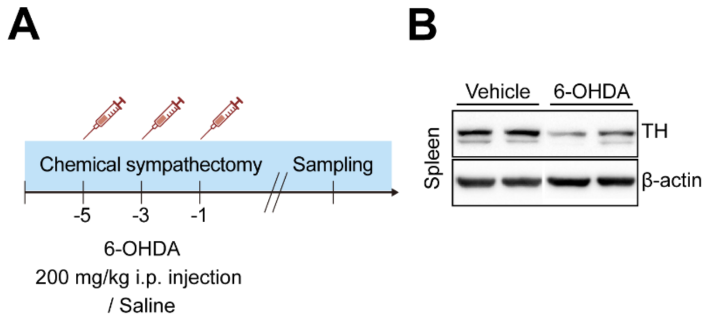


**Supplemental figure 7. The level of TH and NE in the spleen decreased in mice treated with 6-OHDA.**

**(A)** A schematic representation of injection and sampling schedule. **(B, C)** Validation of chemical sympathectomy a day after the injection of 6-OHDA (200mg/kg, *i.p.*). The protein level of TH significantly decreased in spleen of the 6-OHDA injected mice. Data represent mean ± SEM (n=3-4 mice per group); $*p < 0.05$, two-tailed unpaired *t*-test. **(D)** Level of NE also significantly decreased in the spleen of 6-OHDA-treated mice. Data represent mean ± SEM (n=3 mice per group); $**p < 0.01$, two-tailed unpaired *t*-test. 6-OHDA, 6- Hydroxydopamine; *i.p.*, intraperitoneal; TH, Tyrosine hydroxylase; NE, Norepinephrine

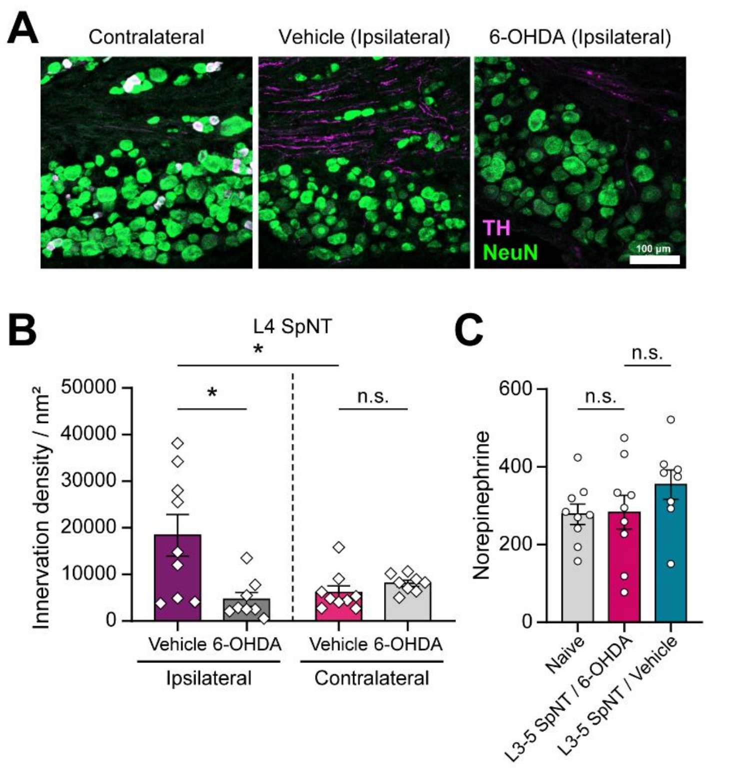


**Supplemental figure 8. The level of TH in the DRG decreased in mice treated with 6-OHDA.**

(**A, B**) Representative immunostaining images and quantitative analysis of TH-IR fibers ($nm^2$) (red) in NeuN+ DRG neurons ($\mu m^2$) (green) in 6-OHDA- vs. vehicle-treated mice at 15 days after L4 SpNT. Data represent mean ± SEM (n=8-9 mice per group, average of 5 sections per mouse); $^*p < 0.05$, two-tailed unpaired *t*-test. (**C**) Quantification of NE levels in L3–L5 DRGs in 6-OHDA- *vs.* vehicle-treated mice at 15 days after L3-5 SpNT. NE levels tended to be lower in 6-OHDA–treated mice compared with vehicle-treated groups, although the difference was statistically not significant. Data represent mean ± SEM (n=8-9 mice per group; pooled samples of two mice per group); two-tailed unpaired *t*-test. 6-OHDA, 6-Hydroxydopamine; TH, Tyrosine hydroxylase; L4 SpNT, L4 Spinal nerve transection; NE, Norepinephrine; n.s., not significant

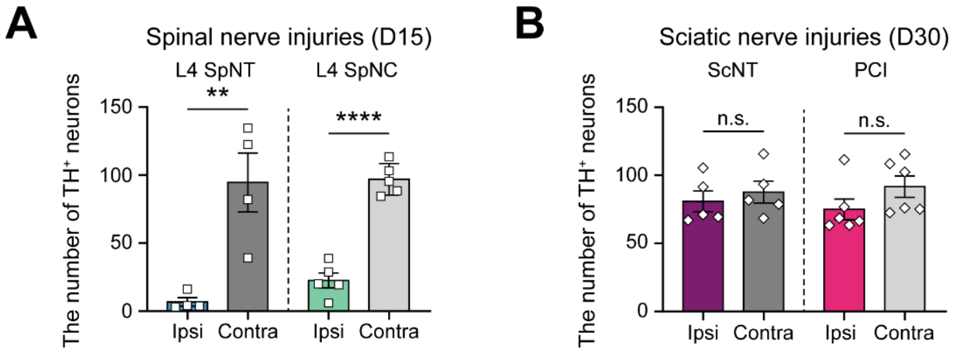


**Supplemental figure 9. The number of TH+ DRG neurons following L4 SpNT, L4 SpNC, ScNT and PCI.**

**(A)** The number of TH+ DRG neuron at 15 days after L4 SpNT and L4 SpNC. The number of TH+ neurons in ipsilateral DRG decreased following both injuries. Data represent mean ± SEM (n=4-5 mice per group, average of 5 sections per mouse); ***p* < 0.01, *****p* < 0.0001, two-tailed unpaired *t*-test. **(B)** The number of TH+ DRG neuron at 30 days after ScNT and PCI. The number of TH+ neurons was not changed in ipsilateral DRG following both injuries. Data represent mean ± SEM (n=4-5 mice per group, average of 5 sections per mouse); two-tailed unpaired *t*-test. TH, Tyrosine hydroxylase; DRG, Dorsal root ganglia; L4 SpNT, L4 Spinal nerve transection; L4 SpNC, L4 Spinal nerve crush; ScNT, Sciatic nerve transection; PCI, Partial sciatic nerve crush injury; n.s., not significant

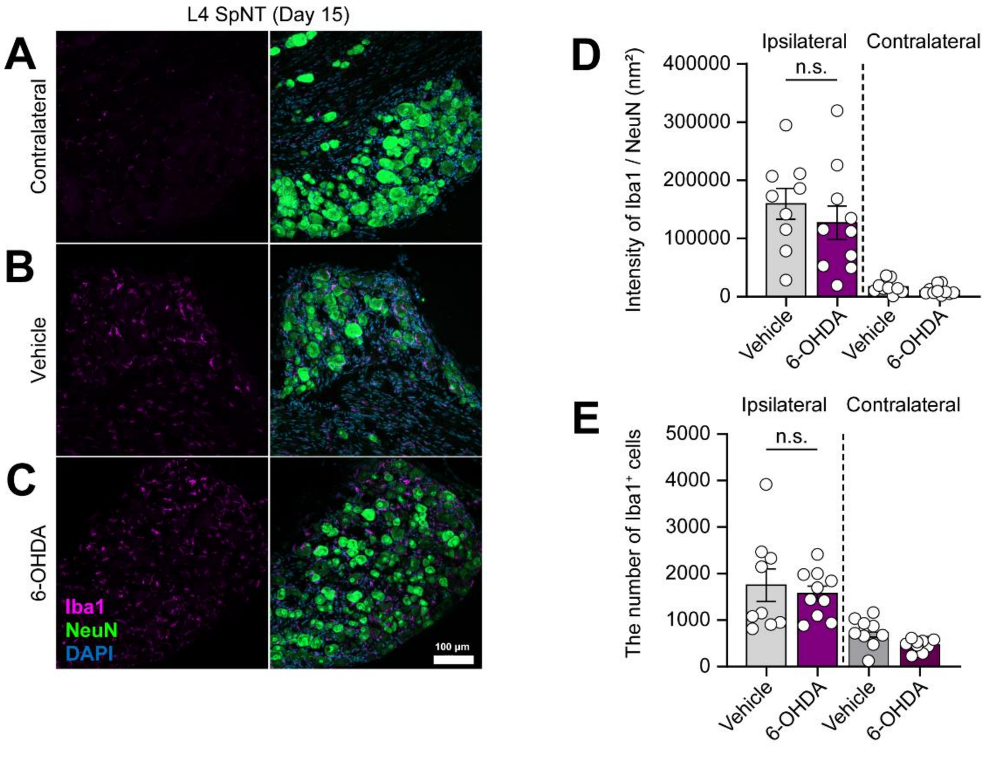


**Supplemental figure 10. The intensity of Iba1+ macrophages is not significantly decreased following 6-OHDA treatment in L4 SpNT mice.**

**(A, B, C)** Representative immunostaining images and quantitative analysis of Iba1-IR macrophages ($nm^2$) (magenta) in NeuN+ DRG neurons ($\mu m^2$) (green) in 6-OHDA- *vs.* vehicle-treated mice at 15 days after L4 SpNT. Data represent mean ± SEM (n=9-10 mice per group, average of 5 sections per mouse); two-tailed unpaired *t*-test. IR, Immunoreactive; 6-OHDA, 6- Hydroxydopamine; L4 SpNT, L4 Spinal nerve transection; n.s., not significant
